# Computational discovery of tissue morphology biomarkers in very long-term survivors with pancreatic ductal adenocarcinoma

**DOI:** 10.1101/207969

**Authors:** Jacob S. Sarnecki, Laura D. Wood, Christopher L. Wolfgang, Ralph H. Hruban, Anirban Maitra, Denis Wirtz, Pei-Hsun Wu

## Abstract

Pancreatic ductal adenocarcinoma (PDAC) is one of the deadliest forms of cancer, with an average 5-year survival rate of only 8%. Within PDAC patients, however, there is a small subset of patients who survive >10 years. Deciphering underlying reasons behind prolonged survival could potentially provide new opportunities to treat PDAC; however, no genomic, transcriptomic, proteomic, or clinical signatures have been found to robustly separate this subset of patients. Digital pathology, in combination with machine learning, provides an opportunity to computationally search for tissue morphology patterns associated with disease outcomes. Here, we developed a computational framework to analyze whole-slide images (WSI) of PDAC patient tissue and identify tissue-morphology signatures for very long term surviving patients. Our results indicate that less tissue morphology heterogeneity is significantly linked to better patient survival and that the extra-tumoral space encodes prognostic information for survival. Based on information from morphological heterogeneity in the tumor and its adjacent area, we established a machine learning model with an AUC of 0.94. Our analysis workflow highlighted a quantitative visual-based tissue phenotype analysis that also allows direct interaction with pathology. This study demonstrates a pathway to accelerate the discovery of undetermined tissue morphology associated with pathogenesis states and prognosis and diagnosis of patients by utilizing new computational approaches.

## Introduction

Pancreatic ductal adenocarcinoma (PDAC) is one of the deadliest cancers, accounting for 7% of all estimated cancer-related deaths in 2016 while only representing 3% of all new cases ^1^. This trend is projected to worsen, with pancreatic cancer projected to account for 4% of all new cancer cases by 2030 but, alarmingly, be responsible for over 10% of patient deaths ^2^. Currently, pancreatic cancer survival rates are very low, with an overall 5-year survival rate of only 8% compared to an overall five-year survival rate of 69% across all cancer subtypes ^3^.

Pancreatic cancer is especially lethal once it becomes metastatic. An estimated 60% of patients with pancreatic cancer are not diagnosed until the cancer is already metastatic, and these patients’ median survival length is less than six months ^4^. However, around 20% of individuals with pancreatic cancer are diagnosed while the cancer is still local and resectable. The five-year survival rate of these patients increases to 20% ^1^, with a median survival around 20 months ^4^. Further, several groups have identified a subset of very long time surviving (VLTS) patients, whose median survival is more than 10 years ^5–10^.

It is unclear if any phenotype can robustly identify this small subgroup of patients with favorable outcomes for patients with more aggressive cancer. Traditional classification metrics, including tumor node metastasis (TNM) classification, do not provide useful information for accurate prognosis of VLTS patients ^6^. Additional work has been done to retrospectively identify clinical parameters that add prognostic value to initial diagnosis. Within these studies, few clinical features, such as perineural invasion or low AJCC grade ^10^, are independently associated with VLTS patient outcomes. Other models using both categorical clinical features and continuous clinical measurements, such as BSA or CA19-9 levels, have achieved similarly poor performance in stratifying patient survival ^11^.

Surprisingly, common genetic mutations do not explain the difference between VLTS patients and pancreatic cancer patients with a poor outcome^12^. However, despite the lack of genetic differences between VLTS and other patients, there exist phenotypic differences in the protein expression patterns ^13^. For example, galectin-1 expression immunohistochemistry scoring can identify VLTS patients with an overall accuracy of 72% ^9^. Interestingly, galectin-1 is only overexpressed in stromal cells within the tumor and is absent in all epithelial adenocarcinoma cells, highlighting the importance of stromal cells in PDAC progression. *In vitro* studies of galectin-1 function find that this protein participates in stromal remodeling ^14,15^ and fibrin degradation^16^. Additional research has identified the tumor microenvironment and stromal cell influences on PDAC progression and prognosis ^17,18^. Notably, genetic analysis of tumor cells and stromal cells separately elucidates specific subtypes that have prognostic value in PDAC ^19^. Recent studies have also identified collagen alignment as a predictive feature for PDAC prognosis ^20^.

Taken together, these results suggest the need for an analytical framework to incorporate information from tumor cells, stromal cells, and their interactions in order to better characterize PDAC. Several *in vitro* ^21–23^ and *in vivo* animal models ^24^ have already been developed to incorporate these tumor-stromal interactions. Yet, the comparability of these models to PDAC in patients is difficult to determine due to lack of quantitative insight from PDAC patient tissue.

Here, we present an image-processing pipeline to extract relevant nuclear, cytoplasmic, and extracellular information from whole-slide images using a computationally-efficient tiling approach ^25–28^. We employ dimensional reduction and clustering techniques to classify each tile according to specific phenotypes and use this clustered information to assess the heterogeneity of the tumor and its surrounding stromal areas. Using this information, we can successfully delineate VLTS patients from those who survive a very short time, suggesting there is exploitable phenotypic information in these whole slide images that could suggest future direction for more basic biological studies. Notably, we find that the inclusion of spatial information outside of the tumor- but in a nearby area- significantly improves model performance and enables increased separation between patients surviving a very long time and those who do not.

## Results

### Visually-aided tissue organization analysis and quantification

Whole slide images of pancreatic tumors from a cohort 29 patients (**Table S1**) including 7 VLTS and 21 STS were first collected. Both VLTS and STS in this cohort have a similar distribution in key disease characteristics, including PNI, TNM, and stage (**Table S1**). However, the majority of VLTS patients have LVI (6 out of 7 patients) in comparison to only approximate half (11 out of 21 patients) STS patients, and STS patients have a larger size of tumor then VLTS when diagnosed. We propose an analytical platform to identify distinct, pathologically-relevant tissue phenotypes to quantify whole slide images (WSI) using the framework presented in **Figure 1a**. Briefly, each/a raw WSI is converted into two nuclei-rich and extracellular matrix-rich areas in the WSI using the previously published nonlinear tissue-component discrimination (NLTD) method ^29^. To minimize the memory loading of the computer, WSIs are computationally partitioned into an array of smaller tissue tiles (500 μm × 500 μm). Within each tile, the tissue morphology and micropatterns were quantified using component intensity and texture features for the quantitative representation of individual tissue morphology. The feature-vectors from each tile are collected from patients, and k-means clustering is used to reduce the complexity of the data and identify characteristic tile morpho-types (TMT). TMTs are representative of subsets of tiles with similar image intensity and texture features. Unsupervised clustering results are visually verified to ensure intra-patient similarity in the TMTs, as demonstrated in **Fig. 1b**. Importantly, the TMTs were further annotated by pathologists to ensure that each TMT contained pathologically meaningful phenotypes. For example, TMTs that encapsulated mainly tissue artifacts, such as tissue folding, or excess blank spaces, such as the edge pieces of tissue, were subsequently excluded from further analysis.

**Figure 1.**
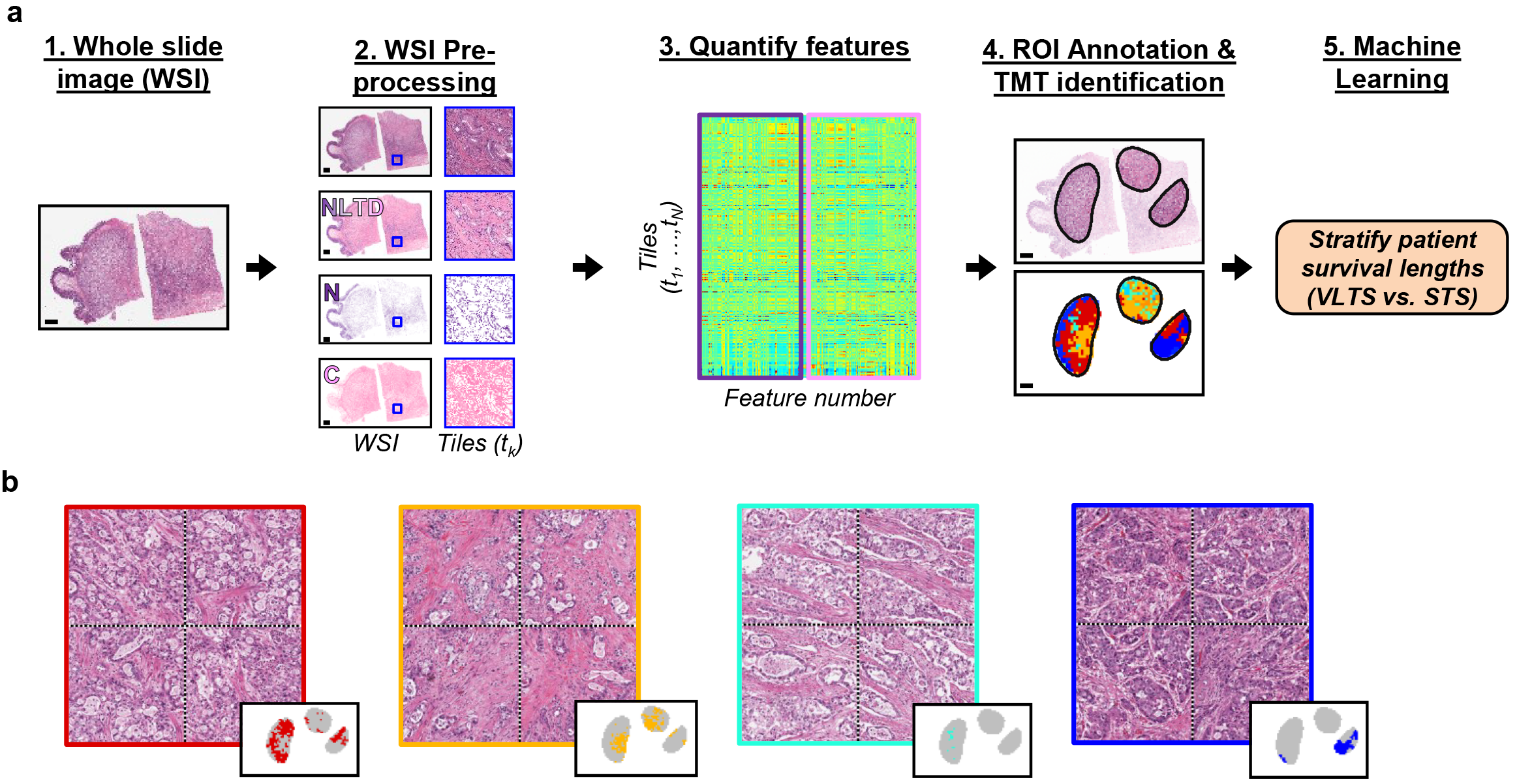
(a) Analysis framework overview for the processing of whole slide images to stratify long-term surviving PDAC patients. Briefly the whole slide image is first transformed into two, grayscale images representing each tissue component, either nuclei-rich regions (N) or ECM/cytoplasmic-rich regions (C) through the NLTD method. After transformation, the image is partitioned into tiles where a series of features is quantified, including intensity and texture features for both tissue components. The feature-set from the tiles are clustered for cancer regions, identified by pathologists, and used to create a multivariate model for PDAC patient stratification. (b) Visual example of clustering results. Each cluster is representative of certain tissue phenotypes that can easily be visualized to create an atlas of clinical features for each cluster. Cluster locations within the whole-slide image can be seen in the inset of each panel.

### Cluster frequency delineates patient survival

We investigated the TMTs that occurred more or less frequently in VLTS patients compared to STS patients (**Fig 2a**). Within both the VLTS and STS patient cohorts, there was significant inter-patient variability in TMT occurrence, with some TMTs occurring in each patient while others occurred in only a few patients. In the VLTS patient cohort, more than half of the tumor regions displayed phenotypes that included fewer than 3 of the 15 identified TMTs. In contrast, only 1 tumor in the STS patient cohort showed such homogeneous phenotype. By averaging TMT occurrences for each patient cohort, the difference in TMT occurrence across VLTS and STS patient cohorts could be further quantified (**Fig. 2b**). The group-averaged TMT occurrence revealed TMT-4 was predominantly VLTS-related and TMTs-7 and 10 were STS-related (**Fig. 2b-c**).

**Figure 2.**
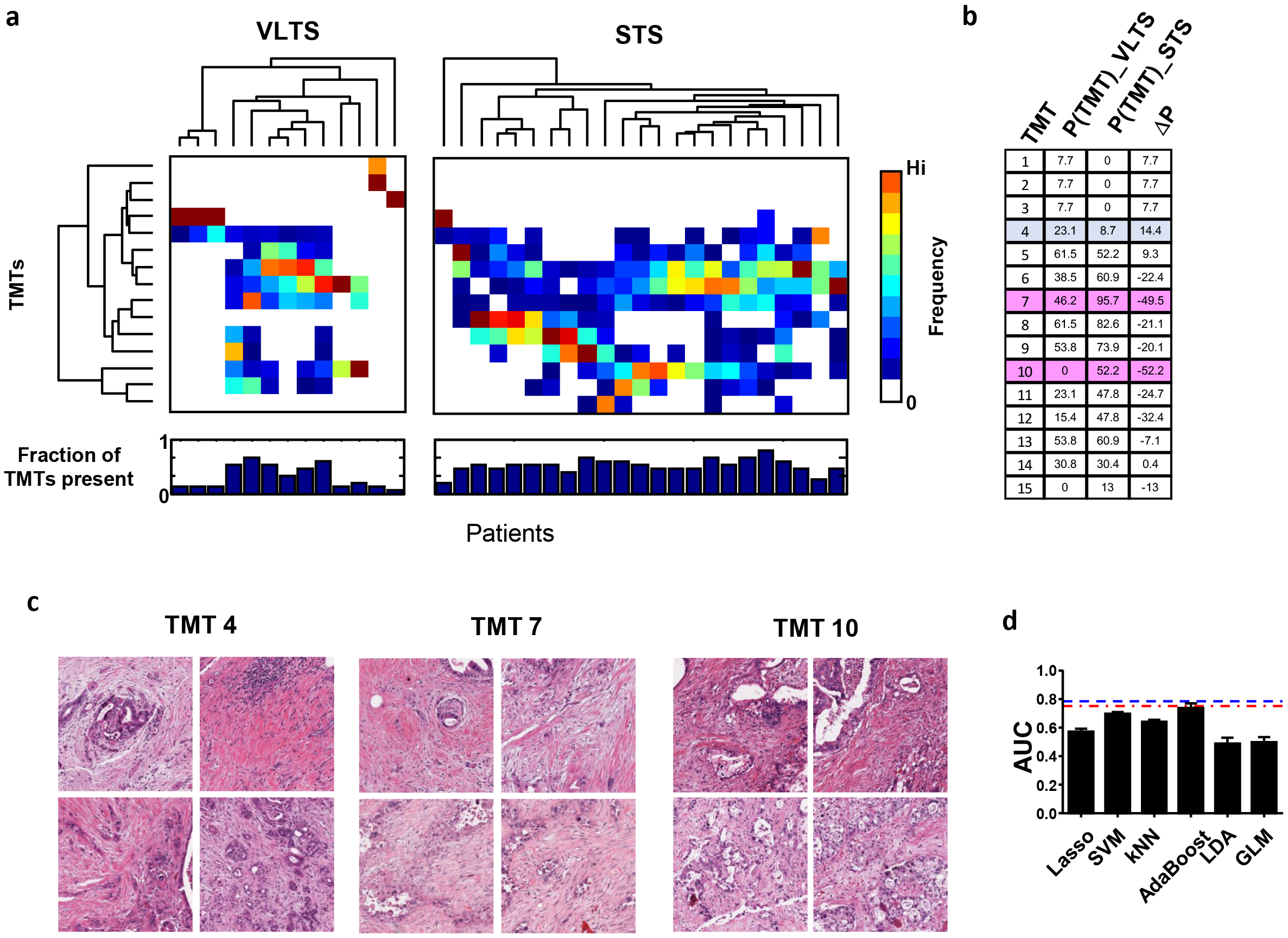
Enriched TMT occurrences in VLTS and STS. (a) Hierarchical clustering of TMT (y-axis) occurrence across VLTS and STS patients (x-axis), with the overall TMT occurrence averaged across patients shown on the far right. Color is representative of TMT occurrence, according to the color scale on the right. Below the clustering results, the fraction of TMTs present in each patient can be seen, with 1 indicating all 15 TMTs are present in the slide. (b) Table show average occurrence of different TMTs for VLTS patients and STS patients and the net difference between this two patient types. (c) Visual demonstration of select TMT phenotypes that are representative of VLTS-heavy phenotype (TMT 4) or STS heavy phentoype (TMTs 7 and 10). (d) Model performance across several multivariate or machine learning models using TMT occurrence information reaches maximal average performance measured by the AUC value. Each bar is the average of 5 independent TMT clustering results, randomly partitioned into training and testing groups 300 times utilizing 3-fold cross-validation, with error bars representing the S.E.M. across the 5 runs.

In addition to visually and qualitatively assessing TMT differences between VLTS and STS patients, we further used TMT occurrence information to build stratification models and assess the predictive power of TMTs quantitatively for patient survival. Patient survival stratification models were built and tested using six different machine learning and multivariate modeling methods (see *Methods* for detailed information) (**Fig. 2d**). To develop the models, the VLTS and STS patient information were randomly partitioned into a training dataset and testing dataset using three-fold cross-validation, where two-thirds of the data were used for training the model, and the remaining one-third was used for testing purposes. The random partitioning was repeated 100 times to ensure robust model performance. Additionally, the cross-validation procedures were repeated for TMT data from five, independent k-means clustering results to account for any variation due to k-means clustering not reaching a global optimum.

Our model performance was varied across the six models, with the AdaBoost model ^30^ showing the highest AUC score of 0.74 and SVM model with radial kernel performing ^31^ next best with an AUC of 0.7. Between the best performing models (LASSO, SVM, k-nearest neighbor, and AdaBoost), there was no statistically significant difference in AUC performance at the 95% confidence level.

### TMT heterogeneity adds predictive value of survival length

In addition to differences in overall TMT frequency between STS and VLTS patient cohorts, there was a striking variation in the number of TMTs present on a patient level, where more than half of the VLTS patients contained fewer than 20% of the identified TMTs (**Fig. 2a**). We reasoned that the number of TMTs present in each patient could be used as an analog for intra-tumoral heterogeneity. For example, if a slide contained only one TMT, it would have a completely homogenous appearance. Conversely, a slide containing each of the fifteen TMTs would have a highly heterogeneous appearance. On average, each VLTS patient contained 31% (95% confidence interval(CI): 18.2%-44.4%) of the 15 total TMTs. In contrast, each STS patient contained 52% (95% CI: 46.3%-57.45%) of the 15 TMTs, representing a statistically significantly higher ratio (*P*=0.001, 95% CI 8.95%-32.25%). This data suggests that the tumor from STS patients is more heterogeneous compared to the VLTS patients since the STS patients have significantly more TMTs present in each slide.

The resultant increase in heterogeneity in STS patients compared to VLTS patients can be seen through representative TMT images of tumors (**Fig. 3a**). In addition to the heterogeneity in TMT frequency, it is apparent that there is also heterogeneity in the spatial distribution of TMTs within the tumor. To more comprehensively quantify the heterogeneity at the patient level, three separate heterogeneity measurement models were used: TMT distribution heterogeneity, gray-level co-occurrence matrix (GLCM) features, and gray-level run length matrix (GLRLM) features. These measures were chosen to account for the different modes of heterogeneity in the WSI, which include the TMT frequency distribution heterogeneity as well as the heterogeneity that arises from the spatial distribution of the TMT in the WSI. An example for each set of calculations can be seen in **Fig. 3b**.

**Figure 3.**
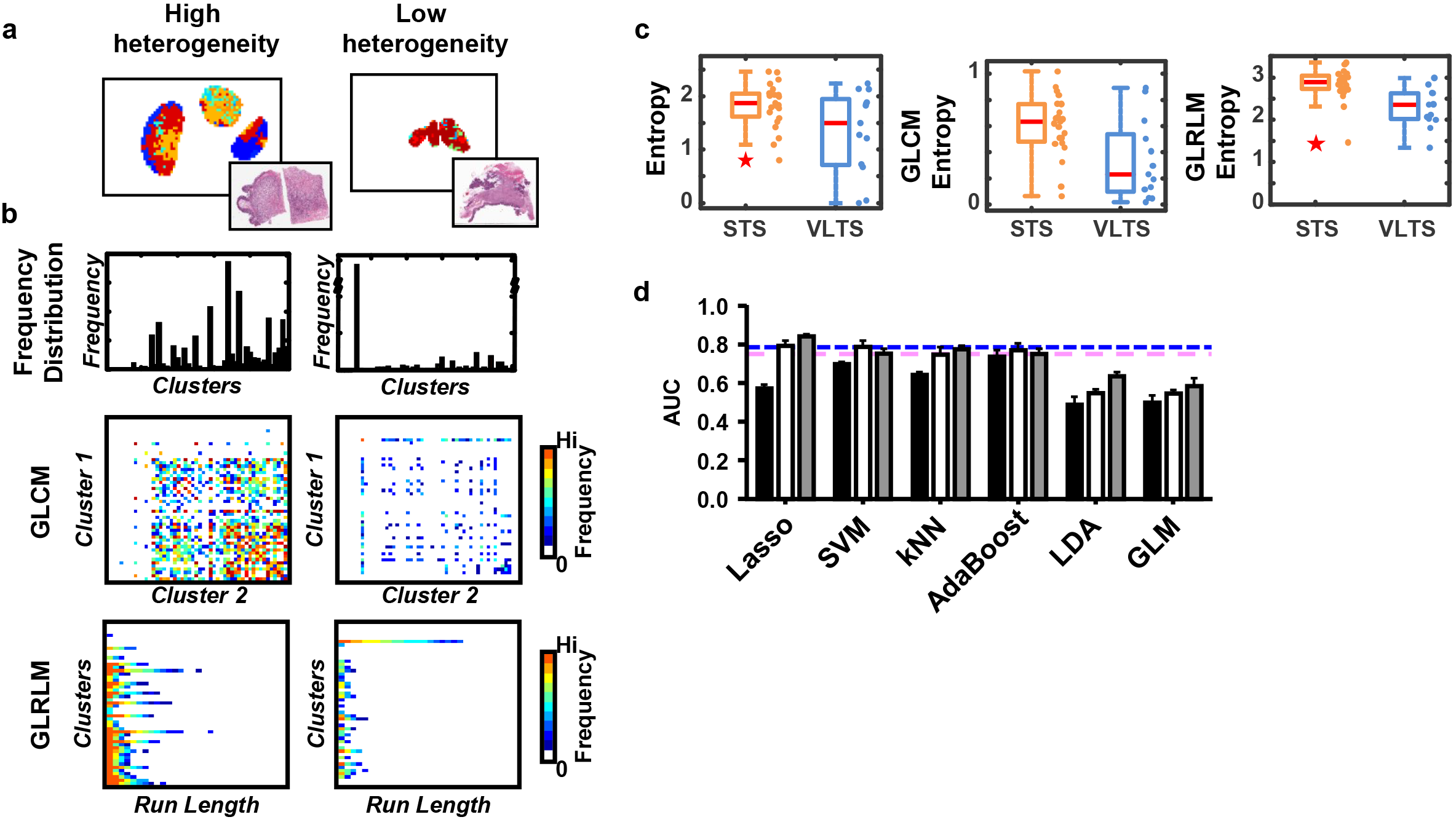
Increased heterogeneity correlated with reduced survival. (a) Representative slides showing a high level (left) and low level (right) of TMT heterogeneity (b) Demonstration of spatial heterogeneity measurements using a high heterogeneity sample (left) and a low heterogeneity sample (right). For each sample, the heterogeneity of the frequency distribution, gray-level cooccurrence matrix (GLCM), and gray-level run-length matrix are derived to quantify overall TMT heterogeneity (c) Heterogeneity measurements from the frequency distribution (left), GLCM (middle), and GLRLM (right) show significant differences between VLTS and STS patients. (d) Each of the three models (TMT frequency, black; spatial heterogeneity, white; combinatorial, gray) was tested using six different multivariate models or machine learning classifiers: LASSO, SVM with radial kernel, k-Nearest Neighbors (kNN), AdaBoost Ensemble modeling, Linear Discriminant Analysis (LDA), and Generalized Linear Model (GLM). Each bar is representative of the average of five separate models built using clustering of tiles within the pathologist-segmented tumor areas with 15 clusters and cross-validated using three-fold cross-validation, repeated with 100 permutations. Error bars are representative of S.E.M. The blue and pink lines represent AUC performance of classifiers built using clinicopathogical features or Galectin-1 immunostaining, respectively.

Briefly, the TMT distribution heterogeneity quantifies the number of TMT present in each slide, as well as the variation in TMT frequencies. The gray-level co-occurrence features measure the heterogeneity between adjacent tiles in local areas of the tumor. The gray-level run-length features examine the persistence of each TMT across adjacent tiles and quantify the short or long run emphasis. The VLTS and STS patient cohorts showed statistically significant differences in heterogeneity, as measured using Shannon’s entropy, for the frequency distribution (*P*=0.028, 95% CI: −0.78, −0.047), GLCM (*P*=0.0064, 95% CI: −0.68, −0.12), and GLRLM (*P*=2.6e−4, 95% CI: −0.85, −0.28). Notably, across all three heterogeneity measures, STS patients exhibited a significantly higher entropy compared to VLTS patients (**Fig. 3c**).

To explore whether a multivariate model based these heterogeneity features could better separate STS from VLTS cohorts, the same six multivariate modeling and machine learning approaches used for the TMT frequency analysis were utilized. These models were trained using a set of heterogeneity measures derived from the three spatial features in **Fig. 3b**. Of these models, the LASSO model had the highest AUC value of 0.793, with the SVM model again having the next best performance with an AUC value of 0.787 (**Fig. 3d**, white bar). For each of the six models, the AUC was furthered improved through incorporating both the TMT occurrence and heterogeneity information (**Fig. 3d**, gray bar). Again, the LASSO model had the highest AUC value of 0.84, with the SVM model having the next best AUC value of 0.78. Importantly, though tumor size for STS patients (3.91±1.2 cm, mean and standard deviation) in our cohorts are slightly larger than VLTS (2.83 ± 1.1 cm), using tumor area as predictor poorly predict the patient survival (AUC of 0.65) and AUC for heterogeneity features based multivariate model does not improve with including tumor (**Fig. S1**). These results suggest that increased tissue heterogeneity in STS vs. VLTS is not due to the tumor area difference.

### Extratumoral area comprises predictive information for patient outcomes

To determine whether the extratumoral area encodes the prognostic value, we tested the model performance when including extratumoral tissues in establishing model (**Fig. 4a**). In addition to the pathologist-identified tumor regions, areas with increased tile radii of 3 tiles (1050 μm), 5 tiles (1850 μm), 10 tiles (3850 μm), and whole slide information were included. For each increased area, k-means clustering and model cross-validation were performed in the same manner as the models built using only the tumor information.

**Figure 4.**
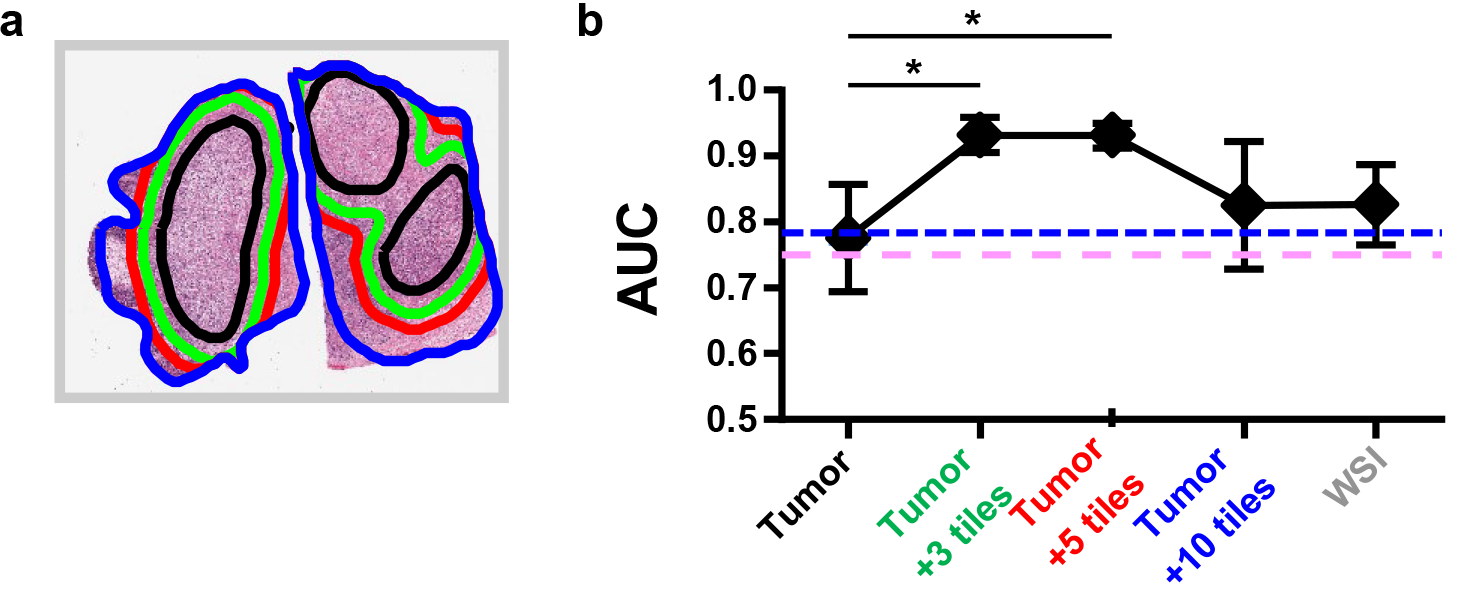
tumor adjacent stromal area encrypt pathological relevant information. Model performance is optimized through the inclusion of peri-tumoral regions in the clustering stage. (a) Pathologist annotations (black) are expanded to include more information at several intervals (green, red, blue) for the clustering process, as well as whole slide information (gray). The inclusion of additional information affects the clustering result, both in cluster phenotypes as well as spatial organization of the clusters, as seen on the right. (b) Using the TMT results from the expanded tumor view a clear dependency is seen with the response of classifier performance to inclusion of additional pathologist information. Importantly, there is an inflection point where the inclusion of more information diminishes model performance at the 5 tile radius (red). The blue and pink lines represent AUC performance of classifiers built using clinicopathogical features or Galectin-1 immunostaining, respectively.

Comparing the performance of the LASSO model across these five regions, we found that the incorporation of extra-tumoral information significantly improved the AUC value at the 3 tile radius (AUC=0.93, p=0.011, CI: −0.28,−0.030) and 5 tile radius (AUC=0.93, p=0.011, CI: −0.28,−0.029) compared to the LASSO result using only tumor information (**Fig. 4b**). Across all six models tested, the LASSO model was found to give the highest overall AUC performance of 0.94 using the 5 tile radius information (**Fig. S2**). To be noted, when including local tissue extended from tumor area in our model, the presence of other tissue types in tissue sections can risk the increased of tissue heterogeneity. Few of our tissue section images show the presence of duodenum, a small intestine tissue. By closed examination by all tissues used in this study, we found an only small fraction of patients in STS (3 out of 21) has positive duodenum tissue close to the tumor and should not substantially contribute to increased tissue heterogeneity in STS patient vs. VLTS when marginally increasing the tissue to analyze based on tumor location.

### Optimal number of TMTs for patient stratification

We examined whether the model performance was sensitive to the number of clusters used in which an arbitrary number less than sample size could be assigned. By definition, increasing the number of clusters used will increase the homogeneity within each cluster, meaning each tile image will have a more similar appearance. The increase in TMT homogeneity would suggest that model performance would continue to improve as more clusters were used. However, we found that as we increased the cluster number, there was an inflection point where the AUC values stopped increasing and began to decrease. To find this optimal cluster number for our approach, we used a series of cluster numbers, ranging from 5 to 50, to test overall performance across different models and the three information types (frequency, spatial heterogeneity, and the combinatorial information).

The results from the two best performing models, LASSO and SVM trained using TMT information from the tumor+5 tile case, are shown in **Fig. 5**. For the LASSO model, the best AUC result was found using 15 clusters for the spatial heterogeneity and combinatorial model. Interestingly, in the LASSO model using frequency information, the best performing cluster number was 50. For SVM, across each of the three information types, the optimal AUC results were achieved using 10 clusters. However, the SVM model performed more robustly with changes in cluster number, with only small differences between AUC values across the cluster numbers tested. Together, these results suggested both under- and over- classification of the tissue morpho-types (TMTs) will decrease the model performance.

**Figure 5.**
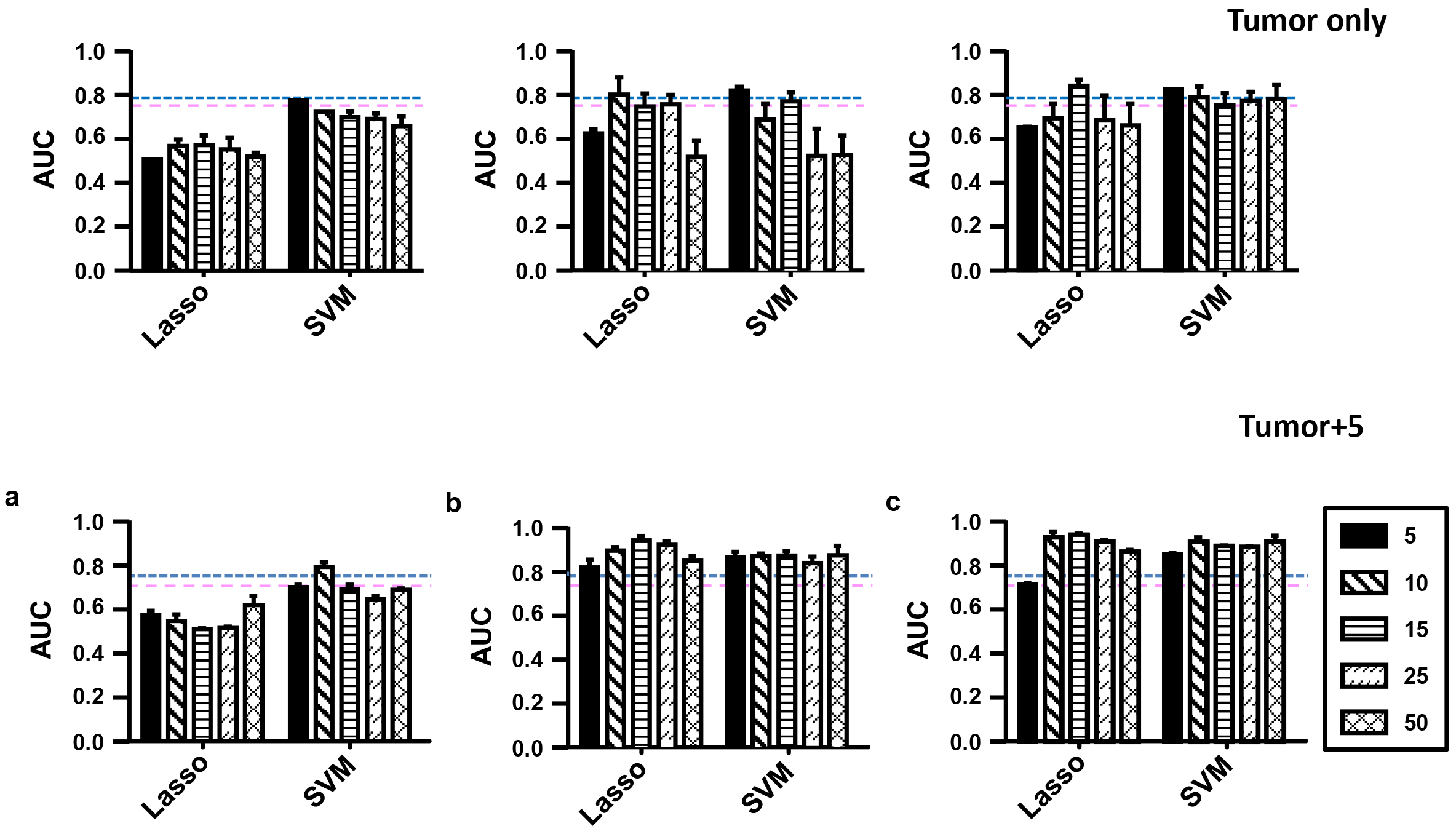
Number of tissue morphs representation classes determines the optimal models performance. (a) LASSO and SVM models predicting VLTS using only frequency did not show any trend with respect to increasing TMT number. (b) The LASSO model using spatial heterogeneity information showed a local maxima at 15 TMTs, while SVM still showed no dependence on TMT number. (c) The LASSO model dependence on TMT information was dampened when frequency information and spatial heterogeneity were combines, but the local maxima still occurred at 15 TMTs. SVM remained TMT independent using the combinatorial model. The blue and pink lines represent AUC performance of classifiers built using clinicopathogical features or Galectin-1 immunostaining, respectively.

## Discussion

To the best of our knowledge, this work represents the first classifier to successfully predict pancreatic cancer patient survival using only information from histopathology images. Previous work in the computational pathology of pancreatic cancer has instead focused on identification of cancerous glands and tissue regions ^32,33^. Moreover, our analysis is the first to extract prognostic information from human, whole-slide tissue images of pancreatic cancer. In this study, we created an image analysis framework that could; (i) automatically normalize tissue staining, (ii) deconstruct whole slide images into smaller tiles, (iii) extract hundreds of image features from each tile, and (iv) use this feature information to facilitate clustering analysis to train multivariate models for the prediction of very long-term survival in pancreatic cancer at time of diagnosis.

Importantly, our pipeline encodes the clustering information back into the original spatial orientation of the image tiles. We found that the spatial heterogeneity information contained the most useful information for modeling VLTS survival. Cluster frequency alone performed significantly worse than spatial heterogeneity information, and the combination of cluster frequency and spatial heterogeneity information only performed marginally better than the spatial heterogeneity features alone (**Fig. 3d**). Though the tiling strategy is a common strategy to reduce WSI information ^26,34,35^, our study suggests that including spatial information can significantly improve disease outcome prediction and patient stratification.

Genomic heterogeneity has previously been identified to contribute to the metastatic potential of pancreatic cancer ^36^. As a result of genomic heterogeneity, it has been hypothesized that tissue morphological heterogeneity might also be associated with PDAC progression ^37^. But tissue morphological heterogeneity has not been systematically studied due to the difficulty to quantify it robustly. Furthermore, a recent study has found that, at the single-cell level, morphological heterogeneity is associated with the metastatic signature of pancreatic cancer ^29^. However, to date, tissue-level heterogeneity has not been robustly quantified in WSIs from patient samples. Our analysis pipeline presents the unique opportunity to quantify tissue morphological heterogeneity utilizing WSIs and, importantly, this quantification of heterogeneity is found to be a strong predictor of patient survival in PDAC and allows for highly robust stratification of VLTS and STS patients.

There is currently no method to robustly identify VLTS patients at the time of diagnosis. Genetic information alone provides no meaningful information ^12^, and both proteomic ^13^ and clinicopathological ^10^ features only offer middling classification performance, with AUC of 0.75 and 0.786, respectively (**Table S3**). While these results suggest better-than-random classification, there is ample room for improvement. Our best model performed with an AUC of 0.94, using information from the tumor and extratumoral region information. Models built using only the clustering information from pathologist segmented tumor regions still perform at a higher level than current state-of-the-art models for predicting VLTS in PDAC patients, achieving an overall AUC of 0.84.

One limitation in this study is the total sample size, which includes 23 STS samples and 13 VLTS samples. This problem is faced by other similar studies, especially in the context of VLTS samples, which are rare ^5,10,13^ With this sample size constraint in mind, we sought to assess the robustness of our model through rigorous cross-validation and re-parameterization. If there were significant outliers in our VLTS sample subset, we would have expected to see larger fluctuations in the model performance across different cross-validation or re-clustering results. We saw very little variation in model performance, as measured by the coefficient of variation for the AUC, in both the cross-validation (μ_Coefficient Variation_=0.16) and re-clustering results (μ_Coefficient Variation_=0.058). Additional analysis with multi-institutional samples will serve to validate the model further, accounting for any variation in tissue staining or preparation that may influence model performance.

In summary, we describe an image analysis workflow that can extract quantitative, texture information from whole-slide images through a tiling approach and recode this information through clustering analysis for spatial analysis. With this pipeline, we demonstrate the utility through the stratification of pancreatic cancer patients who survive for a very long time from those who do not. In this analysis, we find that, while information from pathologist identified tumor areas is predictive, the inclusion of information from regions directly adjacent to the tumor area greatly increases model performance. Furthermore, our system provides pathologists with a platform to visualize meaningful tiles to facilitate better clinical or biological understanding of tissue phenotypes. Overall, our methods present the opportunity to better separate patient subtypes and further analyze tissue phenotypes that are critical to patient stratification.

## Methods

### Patients and sample acquisition

Histo-pathological images were acquired from pathologists at the Johns Hopkins University. The tissue samples were formalin-fixed and paraffin embedded. Tissue sections were fixed for 3 hours in formalin on tissue processor, followed by 1-2 hours of gross room fixation. Paraffin sections were cut at 5μm thickness. Sections were then stained with hematoxylin and eosin and digitized using a DP27 5MP color camera at a resolution of 0.504 μm/px (equivalent to 20X objective). A trained pathologist manually annotated each slide to segment three tumor areas within each tissue. All tissue annotation was done using Imagescope (Aperio) software. Patient samples were carefully selected to account for similar age (μ_VLTS_=66.8±10.1 yrs, μ_VLTS_=66.0±13.2 yrs.) and stage of disease between the VLTS and STS cohorts (**Table S1**).

### Image preprocessing and feature extraction

An overview of the image processing and modeling approach is seen in **Fig. 1a**. First, each whole-slide image was normalized utilizing the Non-Linear Tissue Decomposition (NLTD) method, which has been previously described ^29^. Briefly, the red and blue pixel information is extracted from each image. Using these values, a joint-histogram is created. The local maxima are extracted from the joint-histogram and segmented into lines corresponding to nuclear rich, extracellular/cytoplasmic rich, and red blood cell rich regions. From these segmented lines, a transformation map is created to create separate tissue-type grayscale images from the whole-slide image according to their red-blue color distribution.

Next, to facilitate more efficient processing, whole-slide images were divided into smaller tiles that cover the entire image and reduced four times of spatial resolution (i.e., pixel size of 2μm). The tile size was chosen to emphasize nuclear level features, as well as multi-nuclear organization and larger structures, such as glands. The window was chosen to be a 500 μm square, with 80% overlap between clusters to avoid edge effects. Utilizing this tiling approach, the whole slide images were reduced to between 1500 tiles and 7000 tiles each. Unless otherwise noted, only tiles selected as tumor areas through pathologist segmentation were included in feature extraction and clustering for model testing.

For each tile, a large number of intensity and texture features were computed, similar to a bag of features approach ^38^. The calculated features include intensity summation information, intensity variation, Haralick texture features ^39^, and gray-level run-length features ^40^. The features were calculated on six different image transformations: the nuclear and ECM/cytoplasmic NLTD transformed images with no filter, small bandpass filter, and large bandpass filter. A summary of calculated features by feature and image transformation type is seen in **Table S2**.

### Image tile clustering and frequency evaluation

Once image tile features were calculated for all whole-slide images, the features were combined into a single dataset. The data were normalized using a z-score transformation, and principal component analysis (PCA) was applied to reduce the high-dimensional dataset. The dataset was reduced to the amount of PCA determined-variables to capture 95% of the total variance in the entire dataset. After the data were normalized and reduced, k-means clustering was performed. Moreover, an implementation of the k-means++ algorithm was used in order to further reduce computation times ^41^. Clustering was recursively repeated 500 times to obtain an optimal clustering result and prevent poor clustering from hampering model performance. An example of clustering performance is seen in **Fig. 1b**. Here, the whole slide image from **Fig. 1a** is converted into an image of distinct clusters (**Fig 1a**, **panel 4**), where each color represents an independent cluster. Within this tumor region, it is apparent there are multiple cluster phenotypes present, and examples of four cluster phenotypes are found in **Fig. 1b**, representing the heterogeneity of the tissue and demonstrating the type of clusters that are typically extracted from images using the feature extraction approach.

After clustering, cluster frequency was determined on an individual slide basis as described in **Eq. 1**. For each cluster, ***i***, the frequency was determined by counting the number of tiles, ***t***_***j***_ for each slide, ***j***, belonging to that cluster. This cluster count was normalized by the total number of tiles in each slide, ***N***_***j***_, to give a final frequency, ***f***_***i,j***_.

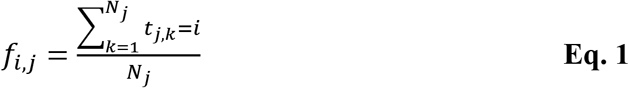

### Tissue heterogeneity measurement

The cluster information was translated back into the image, assigning each tile one value associated with the cluster it represented. This process is visualized in **Fig. 1b**. Once the cluster image is created, cluster distribution and spatial heterogeneity could be quantified. Gray-level heterogeneity features were calculated, including cluster variation statistics, entropy, gray-level co-occurrence matrix features, Haralick texture features, and gray-level run length matrix features (**Fig. 2a**). The gray-level co-occurrence matrix and gray-level run length matrix were calculated for four different angles (0°, 45°, 90°, 135°) and the features were the average from calculation across these four directions. Features were carefully chosen to exclude any features that were influenced by gray-level values (such as mean) since the magnitude of the cluster value did not contain any significant information. Moreover, features were verified such that randomization of cluster magnitude, i.e., changing the value of a cluster from 1 to 10, did not result in a change of output statistics.

### Multivariate modeling and statistical analysis

Multivariate models were constructed using frequency information, spatial information, or a combination of both spatial and frequency information. For spatial information, each feature was normalized across the training and testing datasets using z-score normalization. For frequency information, no further normalization was applied since each cluster frequency was self-normalized across individual slides. Each slide was assigned a binary score according to survival, 1 for very long-term surviving patients and 0 for short-term surviving patients, to prevent overfitting due to the large difference in survival between the two groups. For this binary classification, several models were used: LASSO ^42^, SVM with radial kernel ^31^, k-nearest neighbor classification (kNN) ^43^, an ensemble model using AdaBoost ^30^, linear discriminant analysis (LDA) ^44^, and a generalized linear model (GLM) ^45^.

To test model performance and the robustness of the model, a cross-validation approach was used. Unless otherwise noted, a fold cross-validation approach was utilized, where the dataset was randomly partitioned such that 2/3 of the data used for training and 1/3 of the data used for testing in each model. To ensure model robustness, the random partitioning was repeated 100 times. For each partitioning, the training dataset was used to calculate the mean and standard deviation for z-score normalization of the entire dataset. Additionally, the entire crossvalidation was repeated 5 separate times to account for variation in clustering results, and the average value from these 5 independent cross-validation runs is reported, unless otherwise noted.

For each randomly partitioned testing dataset, the model accuracy, sensitivity, specificity, and area under the curve (AUC) were calculated. AUC was chosen to evaluate model performance. Two-way ANOVA statistics were used to determine performance difference among different classifiers. All reported p-values were corrected for multiple comparisons using Tukey’s method ^46^.

## Acknowledgements

The authors acknowledge funding from the National Cancer Institute (U54CA143868 and R01CA174388)

